# Shifts in ruminant fermentation during inhibition of methanogenesis are reflected in the isotope compositions of volatile fatty acids

**DOI:** 10.1101/2025.10.16.682381

**Authors:** Elliott P. Mueller, Rich Duong, John Eiler, Matthias Hess, Alex Sessions

## Abstract

Ruminant animals are a major source of the potent greenhouse gas methane, but they are also a tractable target for climate solutions. Several strategies have been developed to lower methane emissions from ruminants, including feed additives that inhibit methanogenic archaea. Sustainable solutions must eliminate methane emissions without hampering the microbial fermentation of plant material, which the animal host relies on for carbon and energy. However, current tools cannot directly quantify or characterize the metabolic pathways of in vivo ruminant fermentation. To fill this gap, we developed an electrospray (ESI) Orbitrap mass spectrometry technique to measure the stable isotope ratios (^13^C /^12^C and ^2^H/^1^H) of volatile fatty acids (VFAs) at their natural isotopic abundances directly from rumen fluid. We tested this technique on in vitro incubations of rumen fluid fed three different substrates with and without the additive *Asparagopsis taxiformis*. We found that the isotope composition of VFAs changed and reflected a remodeling of microbial fermentation pathways. Specifically, acetate’s *δ*^13^*C* value increased when methanogens were inhibited, suggesting a lower relative rate of acetate synthesis and a lack of acetogenic activity. Furthermore, the *δ*^2^*H* value of propionate decreased, which may indicate a change in the balance between the two pathways of propionate synthesis toward the less energetic acrylate pathway. Both signals were consistent across feed types. Taken together, our results provide evidence that fermentative metabolism is remodeled during methanogenesis inhibition and decreases relative fluxes through ATP-generating pathways. More broadly, this study demonstrates the utility of ESI-Orbitrap-based isotopic analysis for studying rumen microbial ecology.

**IMPORTANCE:** Slowing methane production from ruminant animals (e.g. cows) is a major target for mitigation of anthropogenic climate change. While strategies that eliminate microorganisms producing methane have been successful, they have cascading impacts on the microbial ecology of the rumen, possibly affecting animal health and productivity.

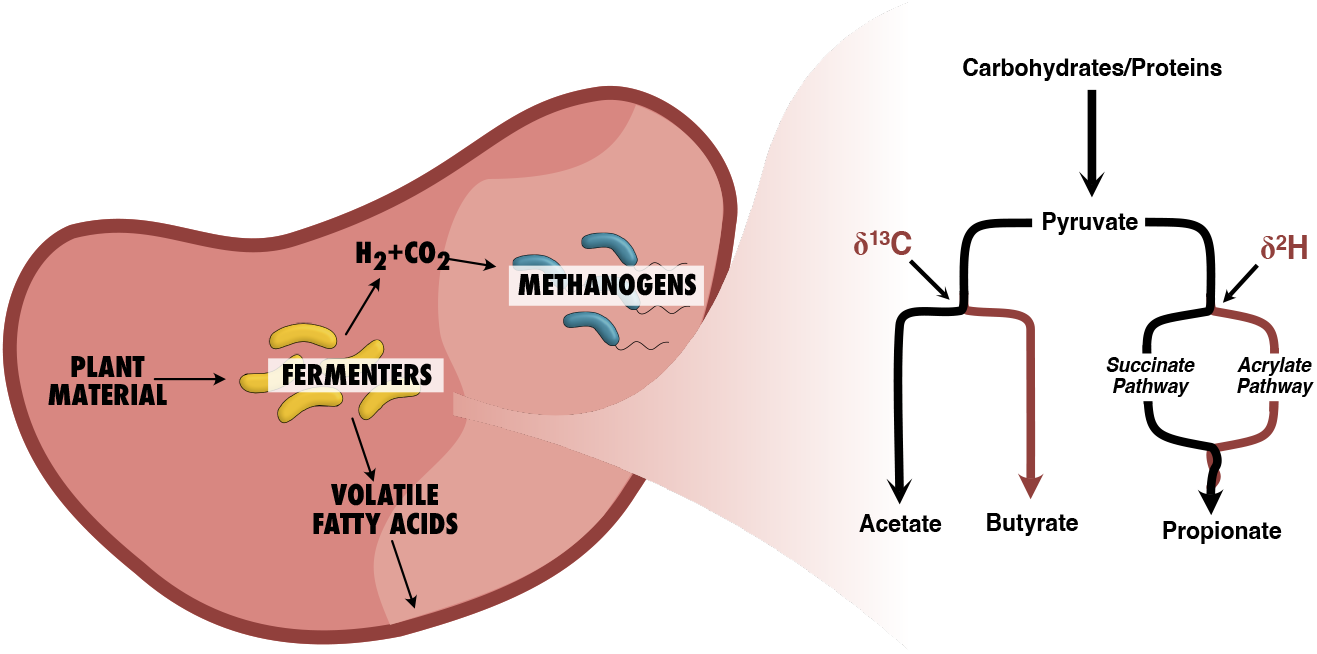

Of particular importance is microbial fermentation, which generates easily digested volatile fatty acids (VFAs) from hard-to-breakdown plant matter. To better understand how fermentation responds to methane mitigation strategies, we measured the isotope composition of VFAs in cow rumen. Our results indicate that fermentation changes pathways when methane production is inhibited to those that generate less energy for the cell. As methane mitigation strategies are developed in the coming decade, isotopic analysis of VFAs may be a useful and accessible contribution to our understanding of rumen microbiology.

## INTRODUCTION

Methane is a potent greenhouse gas responsible for more than 20% of global warming (1). Though lower in concentration than CO_2_, methane traps solar radiation in the Earth’s atmosphere 25-times more efficiently, amplifying its contributions to climate change (2). Livestock agriculture, particularly of ruminant animals (e.g. cows), is the second largest source of anthropogenic methane behind fossil fuels, and is a tractable target for mitigation efforts (3; 4). The anaerobic degradation of fibrous plant materials in the cow rumen (‘enteric fermentation’) creates nearly 500 liters of methane per day per animal (5). With an estimated 1.5 billion cows on the planet, enteric fermentation is responsible for at least 6% of the anthropogenic rise in global temperature (3). In recent years, multiple strategies have been developed to lower methane emissions from cows, including feed-additives. Dosed at low concentrations (<2% w/w), these additives have been shown to drastically decrease methane production in ruminant animals (4; 6). However, we require a system-wide understanding of the response of rumen microbiota to these strategies.

Methane is made in the cow rumen through a cascade of metabolic reactions that breaks down solid plant matter (7). These processes are driven by microorganisms — bacteria, archaea, fungi and protists — densely populating the rumen (∼10^11^ cells/mL) (8). While the physical mechanisms of ruminant fermentation are still an area of active research, the overall structure of carbon flow is well defined. Cellulose — a biopolymer of glucose monomers bound by recalcitrant beta-1,4-linkages — and proteins are first cleaved by exoenzymes excreted from microbial cells. The released monomers are then catabolized by diverse fermentative metabolic pathways. Rather than relying on a terminal electron acceptor, fermenting microbes use the organic substrate as both an electron donor and acceptor, creating CO_2_ and volatile fatty acids (VFAs), respectively (9). The animal host absorbs these VFAs and consumes them for energy (8). Thus, cultivation of fermenting microorganisms in the rumen allows ruminant animals to convert recalcitrant plant matter into bioavailable carbon. However, many fermenting microbes cannot maintain redox balance with VFAs as the only sink of reducing equivalents. They also use water as an electron acceptor, which generates hydrogen gas (H_2_). Methanogenic archaea consume this H_2_ and CO_2_ to make methane. About 12% of the carbon consumed by the bovine host is ultimately released as methane (10). Thus, enteric methane production contributes not only to anthropogenic climate change, but also to carbon loss during digestion for the host. Strategies to eliminate methanogenic archaea from the gut microbiome have become a subject of ongoing research (11; 12).

Feed additives have recently gained traction as an option to mitigate methane emissions from cow rumen fermentation (4; 6; 12; 13). These additives are typically methanogen-specific metabolic inhibitors, namely small halogenated compounds (11). During the fermentation process, they are released into rumen fluid and bind to the active site of methyl-coenzyme M reductase (MCR), a requisite enzyme for methanogenesis (14). By inactivating MCR, these compounds severely limit methane emissions during in vitro incubations of rumen fluid (15). They have also been shown to work in vivo (i.e. in the animal host). The macroalgae *Asparagopsis taxiformis* naturally contains large concentrations of bromoform, an effective MCR inhibitor. When added to feeds, it significantly lowers methane production rates (16; 17; 18).

While additives successfully mitigate methane production, eliminating a major ecological player in the rumen system has consequences for the residual community of fermenting microorganisms. Fermentation and methanogenesis form a syntrophy that expedites plant matter degradation (19). Fermenters feed methanogens with H_2_ and CO_2_ while methanogens continually remove the waste products of fermentation, enabling rapid cellulose decomposition that sustains the animal host. If H_2_ accumulates — which occurs when methanogens are inhibited — fermentation becomes less energetically favorable per mole of glucose consumed, possibly slowing down the total rate of organic degradation (12). Thus, eliminating methanogenic communities could have profound implications for the rumen’s microbial ecology. Indeed, in vitro incubations of rumen fluid with *A. taxiformis* have lower VFA production rates than controls with the same amount of feed (6). The relative abundances of acetate, propionate and butyrate - the major VFAs in the rumen - also change during methanogen inhibition (11; 13). Since these changes are a consequence of the decoupled syntrophy between methanogens and fermenters, they will occur irrespective of the methanogenesis inhibition strategy invoked unless another H_2_ consumer is introduced. Even if microbial communities can be engineered to exclude methanogens during the initial colonization of the rumen, thermodynamic barriers within the cascade of metabolic pathways that breaks down cellulose will still exist. Understanding the response of fermenting communities both in structure and in function is crucial for evaluating the efficacy of engineered solutions and the sustainability of their implementation.

Sustainable strategies would eliminate methane emissions without hampering the fermentation of plant material; however, this is difficult to evaluate in practice. Since multiple fermentative pathways can create the same VFA products, identifying how these pathways change when methanogens are inhibited is challenging even during in vitro incubations. The most common metrics are the total VFA production rate and the relative abundance of these VFAs (11; 13). These can shed light on the response of fermentative organisms; however, their application will become even more limited when strategies are implemented in vivo. In the rumen, organic acids are held in a steady state by the animal host continually consuming VFAs as they are produced (7). In this case, profiles of VFA concentrations will be less informative and their total production rate more difficult to measure. And yet the mechanisms and rates of enteric fermentation are essential for the health and growth of the animal host. Tools are needed that can test the response of fermentation to methane mitigating strategies and can translate to in vivo studies. Here, we developed a novel analytical tool that measures the stable isotope composition of VFAs at their natural abundance in the cow rumen. We demonstrate that these measurements can constrain fermentation pathways during *in vitro* incubations of rumen fluid. We predict that they will also be directly translatable to in vivo studies, where other techniques of tracking fermentation (e.g. VFA profile analysis) are less informative.

Stable isotopes at their natural abundances are useful tools for understanding chemical and biological processes in nature. These tools capitalize on the natural variability of isotopes like ^13^C and ^2^H in biomolecules. The addition of an extra neutron to the atom does not change its broad chemical properties; however, enzymatic reactions often express a kinetic preference for those molecules with lighter (^12^C) isotopes over those containing heavier (^13^C) isotopes (though in certain circumstances the reverse may occur). This kinetic isotope effect (KIE) is realized as a measurable difference between the isotope ratio (^13^C/^12^C) of the reaction’s substrate and its product, known as a ‘fractionation’. The isotope compositions (i.e. *δ*^13^C and *δ*^2^H) of molecules are expressed as a part-per-thousand or ‘permil’ (‰) differences in isotope ratio relative to an internationally recognized standard material.

Since the magnitude of an enzyme’s KIE is dependent on its mechanism, metabolic pathways with different enzymes will express distinct isotope fractionations (20). By measuring the *δ*^13^C or *δ*^2^H values of molecules in nature, it is possible to distinguish metabolic sources of the same molecule. For example, fermentation and reductive acetogenesis both create acetate, the most common VFA in rumen fluid. However, the *δ*^13^C and *δ*^2^H value of acetate are different depending on which metabolism produced it. Fermentation tends to produce acetate with an elevated *δ*^13^C values (0-8‰ enriched relative to the substrate), whereas acetogenesis synthesizes acetate with a strongly deleted (*δ*^13^C <-50‰, *δ*^2^H <-400‰) isotopic signature (21; 22; 23; 24). A recent study also demonstrated that fermenting bacteria express isotopic fractionations between their VFA products that offer information about their different metabolic pathways (25). Differences between VFA *δ*^13^*C* values have even been observed in the rumen (26). Thus, we hypothesized that tandem carbon and hydrogen isotopic analysis of VFAs in the rumen would be a useful constraint on the changing pathways of fermentation and presence of alternative metabolic H_2_ sinks upon in vitro administration of *A. taxiformis*.

Here, we build on a previously developed electrospray ionization (ESI) Orbitrap mass spectrometry (MS) technique for analysis of acetate’s *δ*^13^C and *δ*^2^H values (21). We introduce an innovation to this established technique that enables simultaneous characterization of the isotope compositions of acetate, propionate and butyrate as part of a single measurement — previously, this technique is applied only to pure analytes or one compound of a mixture. (21; 27) To test our hypothesis, we collected fluid and gases from three-day incubations of rumen fluid fed a series of organic substrates with and without *A. taxiformis*. Experiments were performed outside the host *in vitro* to remove the process of VFA consumption. This way, VFA concentration profiles only represented production by the fermentation pathways, enabling us to verify the metabolic shifts identified by isotopic measurements. In addition to the expected decrease in methane emissions, we measured reproducible trends in the isotope compositions of acetate and propionate, which point to shifts away from more energetically efficient pathways when methanogens are inhibited. We predict that the isotopic signals reported *in vitro* here will be captured even in the presence of animal host consumption of VFAs, where VFA concentration profiles become less informative. Taken together, our isotopic analysis of VFAs are useful for constraining fermentation pathways in the rumen that are otherwise invisible.

## MATERIALS AND METHODS

### Rumen Fluid Collection

All animal procedures were performed in accordance with the Institution of Animal Care and Use Committee (IACUC) at the University of California, Davis, under protocol number 19263. Rumen content was collected from a fistulated Holstein cow that was housed at the UC Davis Dairy Research and Teaching Facility Unit. The donor had been fed total mixed ration (TMR). Two liters of rumen fluid and 30 g of rumen solids were collected 90 min after morning feeding. Rumen content was collected via transphonation using a perforated PVC pipe, 500 mL syringe, and Tygon tubing (Saint-Gobain North America, PA, United States). Fluid was strained through a colander into a pre-warmed, vacuum insulated container and transported to the laboratory. Rumen fluid and solids were collected on November 13, 2023. Within 2 hours of collection, the trials had started.

### Rumen Fluid Incubation and Sampling

Rumen incubations were performed in 0.3 L Ankom units (Ankom Technology RF Gas Production System, Macedon, NY, United States). Each unit received 200 mL of a 3:1 mixture of synthetic saliva buffer and rumen fluid. In addition, 2 grams of rumen solids and 2 grams of TMR were added to each unit at the start of the trials. The composition of TMR was 70% alfalfa, 15% dried distillers’ grain and 15% rolled corn. Rumen solids were placed directly in the incubation while TMR was sealed in porous 5 cm × 5 cm concentrate feed bags. For alfalfa and cellulose treatments, 2 grams and 1.5 grams of feed was added, respectively, to the feed bags. *A. taxiformis* was included in the respective feed bags (Ankom, Macedon, NY, United States) at 2% (w/w). The Ankom units were placed and incubated in a shaking water bath (39°C, 40 rpm). Foil gas bags (Restek, United States) were filled with 30 mL of pure nitrogen gas and then connected to the Ankom units. With three different feeds, positive treaments with *A. taxiformis*, and negative controls without *A. taxiformis*, there were six conditions. Every condition included four Ankom replicates resulting in 24 total units run in parallel.

Throughout the experiments, each Ankom unit automatically opened its valve between the gas bag and the incubation headspace when the headspace reached a set pressure. At 24 and 48 hours, each gas bag was replaced with a new one pre-filled with 30 mL of nitrogen gas. At the same time, the Ankom units were rapidly opened, the old feed bag removed, and a fresh one placed in the incubation. Their headspaces were subsequently flushed with nitrogen gas for 30 seconds before closing the units and attaching the new gas bag. When the Ankom units were open, 1 mL of liquid was taken and immediately filter sterilized and placed at -20° C for organic acid analysis.

### CO_2_ and methane concentration measurements

CO_2_ and methane were measured from gas bags every 24 hours using an SRI Gas Chromatograph (8610C, SRI, Torrance, CA, United States) fitted with a 3’ × 1/8” stainless steel Haysep D column. The oven temperature was held at 90° C for 5 min. Carrier gas was high purity nitrogen at a flow rate of 30 mL/min. A 1 mL sample was diluted in 29 mL of pure nitrogen and injected directly onto the column. Calibration curves were developed with Airgas certified methane and CO_2_ standard (Airgas, United States).

### CO_2_ and methane isotopic analysis

The carbon isotope composition of CO_2_ and methane were measured on a gas chromatograph isotope ratio mass spectrometer (GC-IRMS). A Thermo Scientific Trace 1310 gas chromatograph with a GC Isolink II was coupled to a 253 plus 10 kV IRMS. Injection volumes varied from 10 to 100 uL of gas samples through a 100 *µ*L gas-tight syringe. The inlet temperature was held at 30° C. The injection volume and injector inlet split flow values were changed to match the heights of CO_2_ and methane sample peaks with the CO_2_ reference gas peak height of 7 volts. Typically, CO_2_ concentrations were several-fold higher than methane concentrations. A split ratio of 40 was used for CO_2_ measurements, while a split ratio of 3 was used for methane, meaning the sample was injected on two separate occasions to measure each of the two analytes. Carrier gas flow rates were held at 2 mL/min. Using an Agilent PORAPLOT-Q 25 meter column, CO_2_ and methane were sufficiently separated in a 6 minute, isothermal GC run held at 30° C. The *δ*^13^C value of both analytes was anchored to the VPDB scale using a tank of pure CO_2_ with known carbon isotope composition (-12.04‰). To verify the accuracy of the measurement, purified methane of known carbon isotope composition (-42.9‰, (28)) diluted to similar concentrations as the samples (1% for split ratio 3 and 10% for split ratio 40). This standard was run every three samples to monitor instrument performance. Deviations between the measured and known *δ*^13^*C* value of the methane standards were used to correct samples. These deviations did not exceed 2‰. Since biological quadruplicates were available, analytical replicates for each sample were not run. Instead biological reproducibility served as the uncertainty, which is represented in the box plot of Figures 1 and 2.

**Figure 1.**
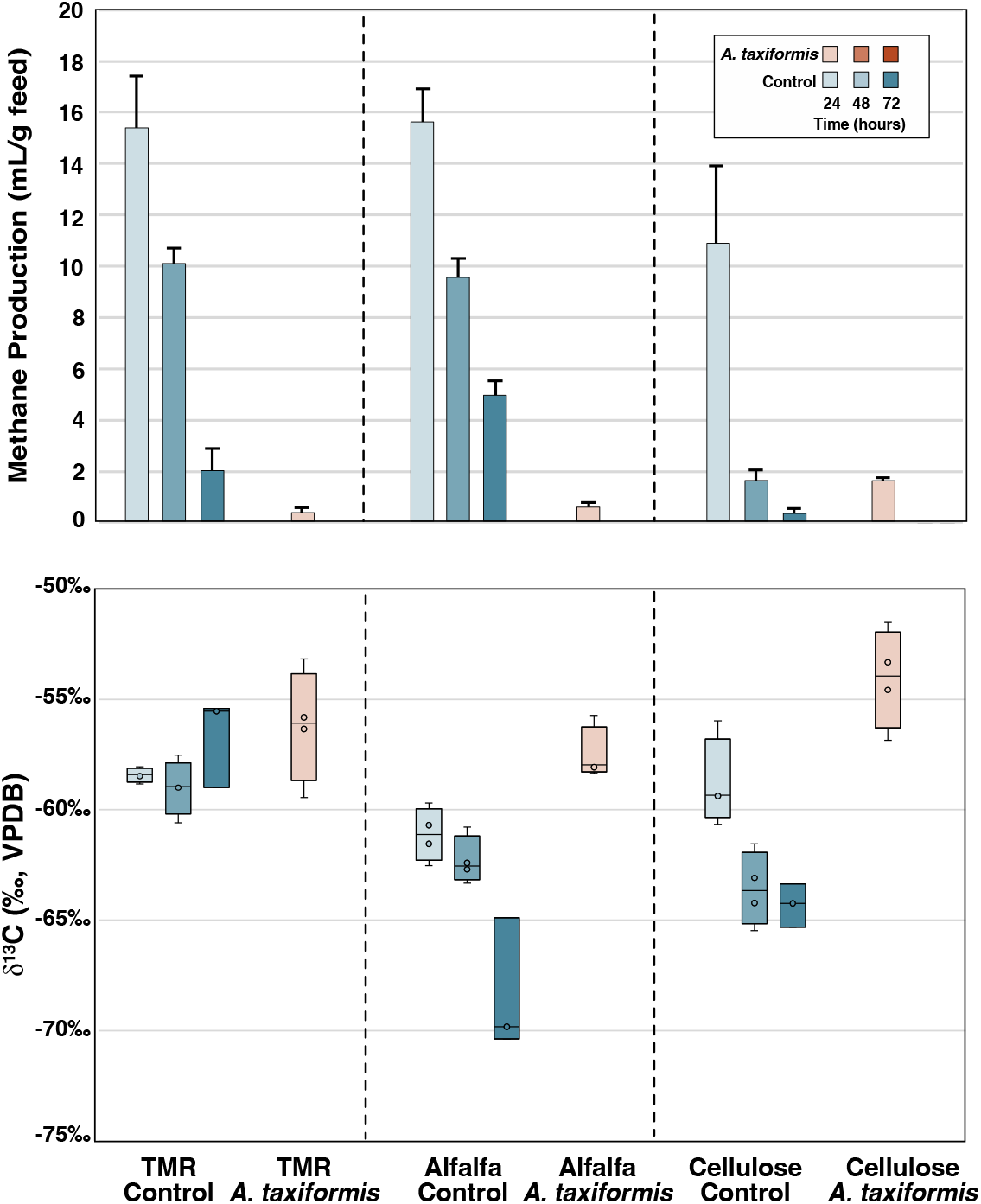
Methane emission rates in the rumen incubations and methane carbon isotope composition over three days. Methane emissions decreased by >90% with the addition of *A. taxiformis* . Blue bars and boxes indicate negative controls without *A. taxiformis* while red boxes indicate a position treatment with *A. taxiformis* .

**Figure 2.**
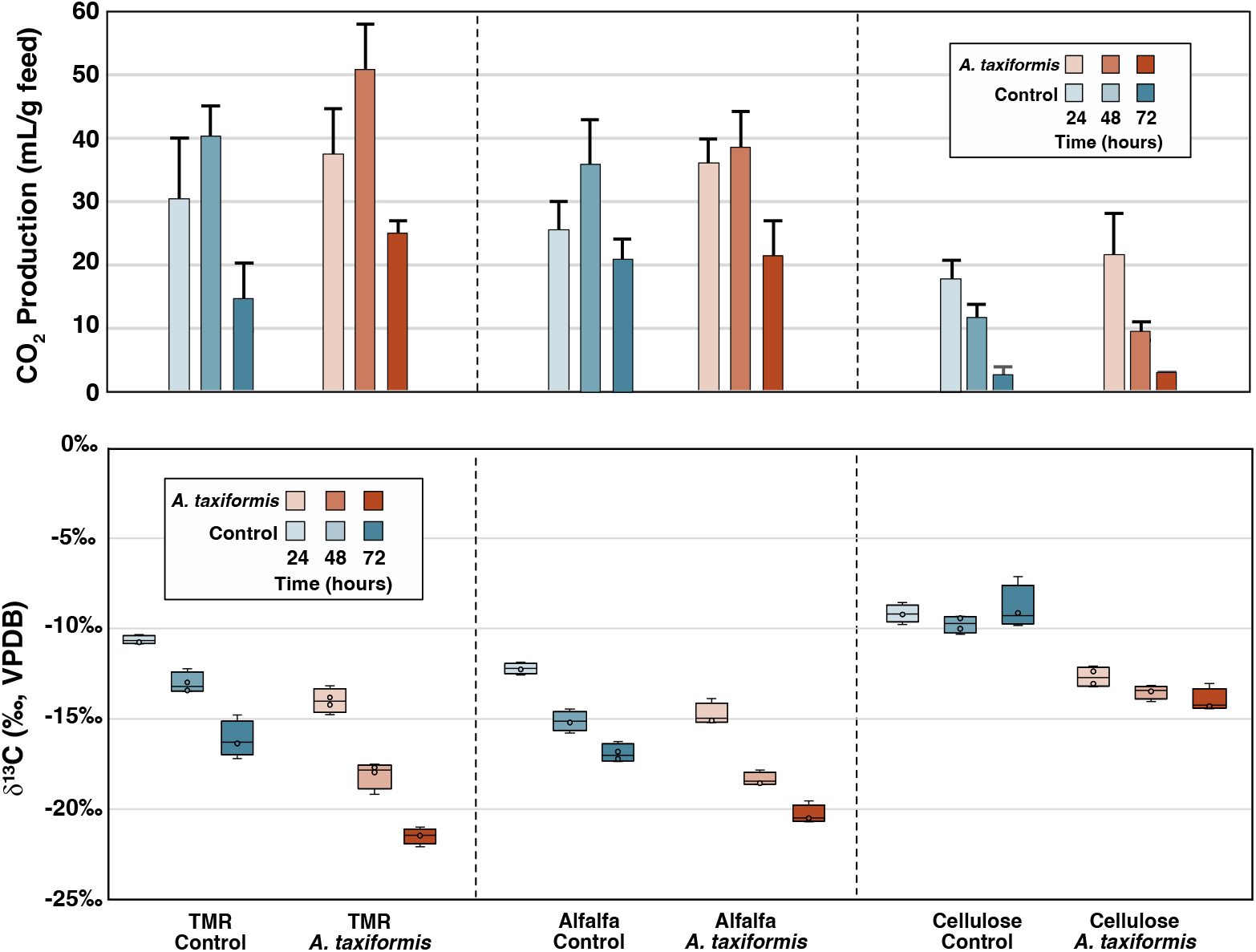
Carbon dioxide production rates in the rumen incubations and its carbon isotope composition over three days. Carbon dioxide production increased slightly upon addition of *A. taxiformis*, while its *δ*^13^*C* balue decreased by ∼5‰ between incubations with (red) and without (blue) *A. taxiformis* . The CO_2_ *δ*^13^*C* values decreased with time when ALF and TMR were used as feed, but not CEL.

### VFA concentration measurements

VFA concentrations in rumen samples were measured using a Hewlett Packard Series 1100 high performance liquid chromatograph (HPLC) coupled to refractive index detector (RID). VFAs were separated on an Aminex HPX-87H (300 x 7.8 mm) column with an isocratic 8 mN sulfuric acid mobile phase. Peak areas were converted to concentrations with an external calibration curve of acetate, propionate, and butyrate from 0.1 to 50 mM. Before injection, all rumen samples were filter-sterilized and diluted 10-fold in 8 mN sulfuric acid. Injection volumes for sampels and standards were 10 *µ*L. Blanks of MilliQ water were run every 3-5 samples to monitor sample carryover, which was not measurable. Calibration curves were rerun at the beginning of every day of the instrument session.

### VFA isotopic measurements

The compound-specific isotope composition (*δ*^2^H and *δ*^13^C values) of acetate, propionate and butyrate were measured on a heated electrospray ionization (HESI) Orbitrap mass spectrometer (MS) (Thermo Scientific) attached to a Vanquish HPLC (Thermo Scientific) with a split-sampler and 100 *µ*L sample loop. No LC column was used for these analyses. Each injection, 100 *µ*L of sample was pulled through the sample loop and then injected into the 5 *µ*L/min flow of LC-MS grade methanol (Fisher chemical, Optima), which carried the sample directly into the Orbitrap MS. After 19 minutes of measurements (2 minutes of dead volume followed by 17 minutes of sample analysis), the flow rate was increased to 35*µ*L/min for two minutes to flush out the remaining sample and then decreased to 5*µ*L/min again to prepare for the next sample. The ESI ionization parameters can be found in Table S1. This 19 minutes was split into three segments, each focusing the quadrupole and Orbitrap on one of the three organic acids. The raw mass spectra were converted to ion counts using the Makarov equation. The ion intensities from mass spectra were converted to counts (N) by applying an empirical factor (C_N_ = 4.4) derived by Makarov and Denisov (2009) and the charge of the ion (z). (29; 30).

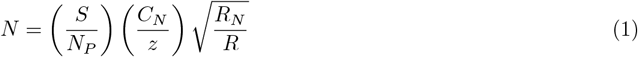

As in (21), the isotope ratios of each organic acid were then calculated from the ion counts and further converted to the delta notation on the VPDB and VSMOW scales. Only data between 3 and 20 minutes were considered for isotope ratio analyses, as this represents the period during which the ion current was stable and in a plateau. On either side of this period, the ion current was either increasing as sample infused into the instrument or was dropping as the last of the sample entered the mass spectrometer. Furthermore, only the periods designated for a given organic acid were used to calculate that organic acid’s isotope composition (e.g. Acetate: 3 to 11 minutes; propionate: 11-17 minutes, butyrate: 17-20 minutes). Each biological replicate was run once and was bracketed by a standard that matched each organic acid’s ion current intensity within 30%.

### Propionate and Butyrate Standards

To anchor the measured ^13^C and ^2^H isotope ratios on the international reference frames Vienna Pee Dee Belemite (VPDB) and Vienna Standard Mean Ocean Water (VSMOW), standards of known *δ*^13^C and *δ*^2^H values were required. Sodium propionate (>99%) and sodium butyrate (>99%) was obtained from commercial sources to create this standard. Its synthetic or biological origins are unknown. Stock solutions (2M) were made up in deionized water (Milli-Q). Batches of working 2M standard stock solutions in MilliQ water at were flash frozen in liquid nitrogen. To ensure homogeneity, stocks were kept as frozen aqueous solutions. Aliquots from these stocks were taken by thawing them at room temperature, inverting the vials to homogenize, aliquoting and immediately re-freezing the stocks. Each standard was measured via Elementary Analyzer (EA) coupled to an Isotope Ratio Mass Spectrometer (IRMS) to independently determine their isotopic compositions relative to the VSMOW and VPDB reference materials. Carbon isotope compositions were measured with combustion EA-IRMS. Samples were calibrated against glycine (-45.7‰) and Urea (-27.8‰) standard. Hydrogen isotope compositions were measured with thermal conversion EA-IRMS using an elemental chromium catalyst as discussed by (31; 21). Samples were calibrated to the VSMOW scale by analyzing USGS77 (polyethylene powder) and C36 n-alkane #2 provided by Arndt Schimmelman (Indiana University). The sodium propionate and sodium butyrate standards are referred to as ProA and ButA. ProA and ButA had *δ*^2^H values of -110‰ (± 1.5‰) and -113‰ (± 1‰), respectively. Their *δ*^13^C values were -34.3‰ (± 0.1‰) and -30.4‰ (± 0.1‰), respectively.

### Feed bulk isotopic analysis

The carbon isotope composition of TMR, alfalfa and cellulose were measured using an Thermo Scientific elemental analyzer (EA) with a ConFlo system coupled to a Delta V IRMS. A total of 5 *µ*g of carbon was weighed out into tin capsules. The EA combusts the biomass and then purifies CO_2_, which is transferred to the IRMS. The CO_2_ ion peak height was similar to the CO_2_ reference gas (-12.04‰). Carbon isotope compositions were corrected against external standards of glucose and glycine with known *δ*^13^C values of -11 and -45.7‰, respectively.

### Metabolic Equations

The balanced equation for the conversion of glucose to pyruvate with H_2_ representing electrons where Pi is orthophosphate below were used to estimate reactions in the rumen. ADP and ATP are adenosine diphosphate, ATP is adenosine triphosphate. All organic acids are assumed to be in their conjugate base at physiological pH.

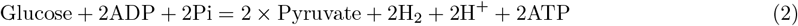

The balanced equation for the conversion of pyruvate to acetate, propionate, and butyrate.

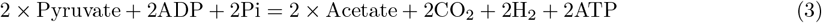

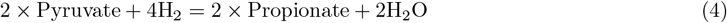

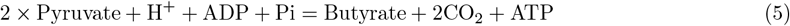

## RESULTS

### Gas Production and Isotope Composition

#### Methane

Cow rumen fluid was incubated with feed bags containing three different organic substrates: total mixed ration (TMR), alfalfa (ALF), and cellulose (CEL). Over the course of three days, the total methane production rates (mL/g feed) in TMR and ALF controls that did not contain *A. taxiformis* were similar, decreasing from 15 mL/g to 2-5 mL/g. With CEL, the decrease also occurred, though the total methane production rates were lower, starting only at 10 mL/g and decreasing to near zero (Figure 1). As a purely fibrous material with no easily digestible proteins or starches, CEL condition was expected to have lower overall gas production. The *δ*^13^C of methane did not vary at the 24 hr time point, even with the addition of *A. taxiformis*. However, when fed ALF and CEL, methane’s *δ*^13^C value shifted to ∼5‰ lower values over 72 hours. This was only observed in negative controls. When *A. taxiformis* was added the incubations, methane was too low in abundance to measure for concentration or isotope composition after 24 hours (Figure 1).

#### Carbon dioxide

Carbon dioxide production rates were also lower in CEL relative to TMR and ALF. However, the trend of decreasing CO_2_ production over time was not as clear with TMR and ALF as compared with the strong decrease in CEL incubations (Figure 2). For each substrate, CO_2_ production rates between incubations with and without *A. taxiformis* were similar. We note that CO_2_ concentrations in the gas bags collected at 24 hour and 48 hour timepoints exceeded the operational range of the instrument. More apparent was the temporal trend in CO_2_ carbon isotope composition. Over three days, the *δ*^13^C of CO_2_ decreased by 5-10 ‰ in every incubation with TMR or ALF. However, the positive and negative treatments (i.e., with and without *A. taxiformis*) were also offset from one another. In TMR and ALF incubations, those incubations with *A. taxiformis* were consistently 2-5‰ lower in *δ*^13^*C* than the same time point in negative controls (Figure 2). When the incubations were fed CEL, there was no temporal change in *δ*^13^C values of CO_2;_ nevertheless, the isotope compositions were offset by 2-5‰ between incubations with and without *A. taxiformis*. CO_2_ produced from those incubations with methane inhibited were more ^13^C-depleted, presumably due to the lack of a consumption reaction that would otherwise leave the residual pool of CO_2_ ^13^C-enriched.

### VFA Concentration and Isotope Composition

The total production of VFAs decreased when methanogenesis was inhibited by the addition of *A. taxiformis*. When fed TMR, ALF, and CEL, the net production of VFAs in negative controls (final minus initial) was 64, 77, and 38 mM VFAs respectively. In positive treatments, net production of VFAs in TMR, ALF, and CEL incubations was 39, 57, and 25 mM, respectively. Error bars in Figure 3 indicate the standard error of the mean (SEM) of quadruplicate Ankom units. Overall, the errors were less than 5% of the mean for most conditions. CEL had the least change in VFA production between negative and positive treatments with acetate being the only VFA that decreased outside of error. This suggested that fermentation did not slow down in the CEL incubation as much as it did in the TMR and ALF incubations when methanogenesis was inhibited. The total VFA production was highest when ALF was added to the incubations. Since TMR contains more fibrous materials than pure ALF, the ratio of easily degraded starches and proteins to fibers in the ALF would have been higher than TMR. This also aligns with CEL having the slowest fermentation rates, since it is entirely fibrous.

**Figure 3.**
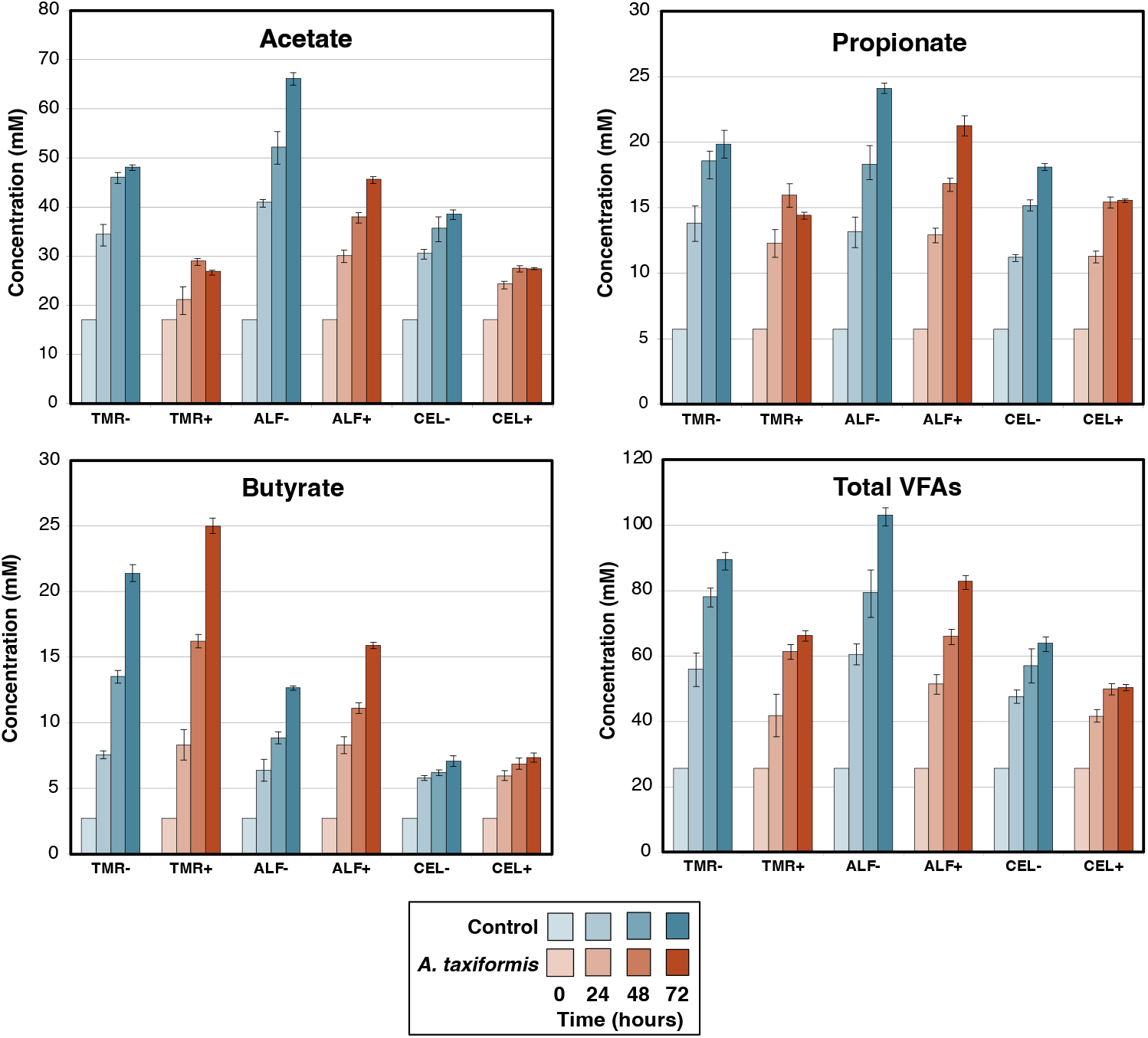
VFA concentrations with time over three day rumen incubations. Acetate and propionate production decreased when *A. taxiformis* was added, while butyrate production increased. These trends persisted across all three feeds. Abbreviations: ALF, alfalfa; CEL, cellulose; TMR, total mixed ration.

The carbon isotope compositions of the VFAs had a consistent inverse relationship with carbon chain length. Acetate (C_2_), propionate (C_3_) and butyrate (C_4_) had successively lower *δ*^13^C values in the initial rumen fluid. These trends were less consistent as the incubations proceeded, though the overall pattern of *δ*^13^C_Acetate_ >*δ*^13^C_Propionate_ >*δ*^13^C_Butyrate_ persisted. In negative controls, the hydrogen isotope composition of the three organic acids were remarkably similar, all around -210‰. Overall, the biological reproducibility of the carbon and hydrogen isotope ratios was notably high with all four incubation replicates falling within analytical error (0.5‰ for carbon, 5‰ for hydrogen) under every condition. Error bars in Figure 4 represent the SEM of these replicates and are mostly encapsulated by the data markers. This level of reproducibility suggests that the isotope compositions of the VFAs are a robust signature of carbon flow. Below, we highlight key results from each VFA in terms of concentration and isotope composition.

**Figure 4.**
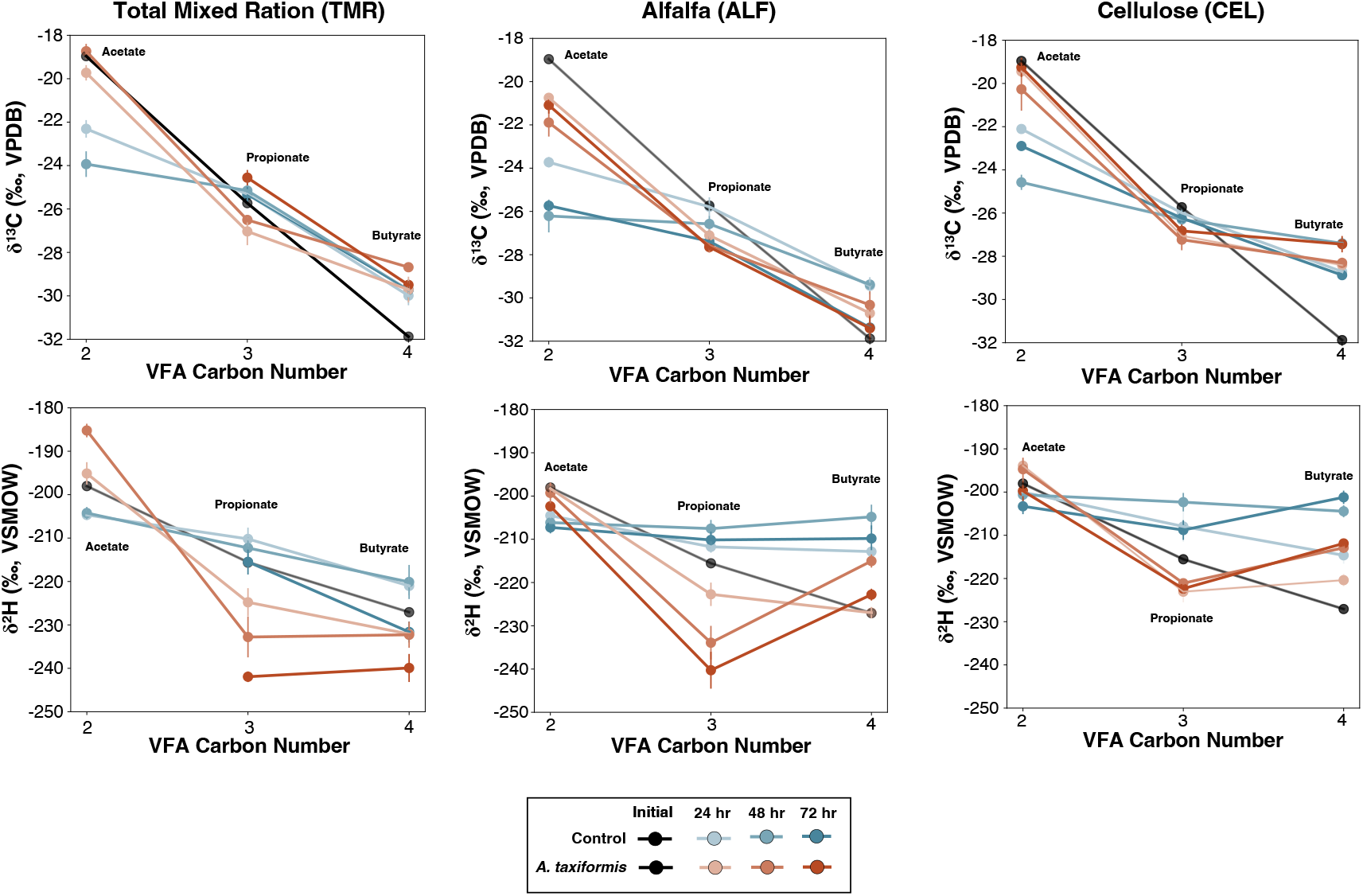
The carbon and hydrogen isotope composition of VFAs over three days of rumen incubations plotted against carbon number, with two, three and four representing acetate, propionate and butyrate respectively. These trends are plotted for incubations fed TMR (A), ALF (B) and CEL (C). Black, red and blue lines represent initial conditions, positive treatments with *A. taxiformis*, and negative controls without *A. taxiformis*, respectively. Shading of the color indicates the time with darker hues representing later timepoints. Error bars represent standard error on isotopic compositions from four incubation replicates

#### Acetate

Acetate concentrations changed with time, the type of feed, and the presence or absence of *A. taxiformis* additive. In all negative controls (no *A. taxiformis*), acetate concentrations rose monotonically from 17mM to 50 mM, 65 mM, and 40 mM with TMR, ALF, and CEL respectively. While there was still a rise in concentration over time, the TMR, ALF, and CEL incubations only accumulated acetate to 28 mM, 45 mM and 28 mM, respectively, when *A. taxiformis* was added (Figure 3). The decrease in acetate production is likely due to high H_2_ partial pressures that makes acetate production less thermodynamically favorable; however, this hypothesis is speculative given that H_2_ partial pressure was not measured.

Acetate became more ^13^C enriched when methane was inhibited (Figure 4). Initially, acetate had a *δ*^13^C value of -19‰, about 8‰ higher than the TMR (-27‰). In negative controls, acetate was more ^13^C-depleted with a *δ*^13^C value of -22 to -26‰, but did not vary systematically with time. However, when *A. taxiformis* was added, acetate was more ^13^C-enriched, on average (between -22‰ and -19‰). We note that the initial condition likely represents a combination of isotope fractionations in VFA synthesis and VFA consumption, while the experiments only capture the former. The hydrogen isotope composition of acetate was steady throughout the experiments, shifting between -190 and -210‰, similar to the initial condition.

#### Propionate

Under all conditions, propionate accumulated in the incubations with small offsets between negative controls and positive treatments with A. taxiform. Initial propionate concentrations were ∼5 mM and rose to 24 mM, 18 mM and 20 mM in negative controls of TMR, ALF, and CEL, respectively. When *A. taxiformis* was added, total propionate production decreased by approximately 20% relative to negative controls. In the CEL and TMR conditions, the last 24 hours of the experiment propionate production slowed, while in the ALF condition rate of propionate production was consistent (Figure 3).

Propionate (Figure 4) had a consistently lower *δ*^13^C value (∼-26‰) compared to acetate. Its carbon isotope composition did not vary over time, with different feeds, or after methane inhibition. However, the *δ*^2^H value of propionate changed over time in TMR and ALF conditions when *A. taxiformis* was added to the incubations (Figure 4A and B). Propionate’s *δ*^2^H value shifted by -30‰ over the course of the experiments (Figure 5). However, this trend had a smaller slope when cellulose was the feed source, changing by only -10‰ when *A. taxiformis* was added.

**Figure 5.**
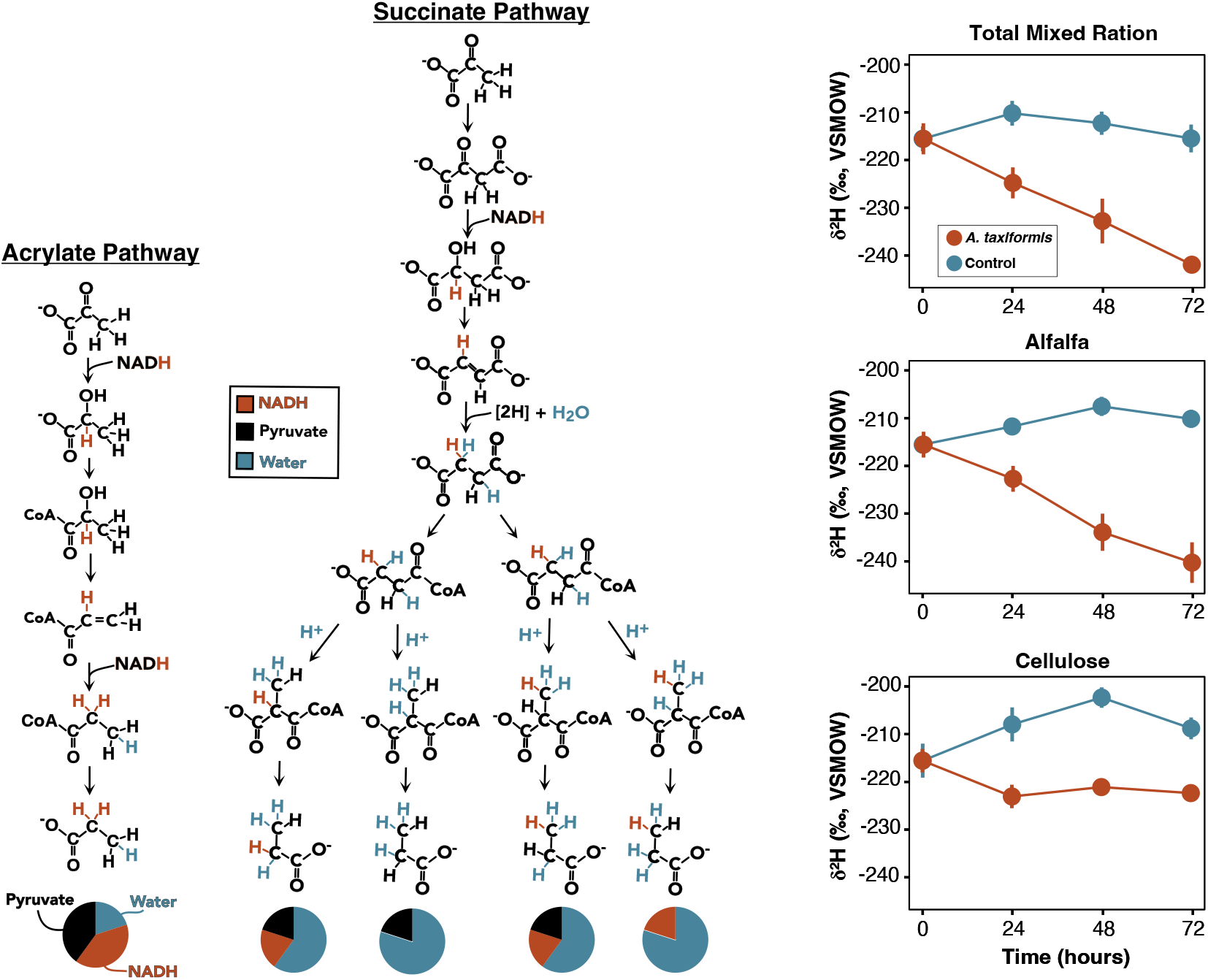
Propionate *δ*^2^*H* values indicate a shift in synthesis mechanism. On the left are the two biosynthetic pathways for generating propionate from pyruvate. The three sources of hydrogen atoms — pyruvate, NADH, and water — are color coded black, red, and blue, respectively. Branch points in the succinate pathway represent reactions involving molecular symmetry in which two distinct isotopologues are possible products of the same reaction. On the right, time-varying *δ*^2^*H* values of propionate when incubations were fed different feed types with (red) or without (blue) *A. taxiformis* .

#### Butyrate

The production rates of butyrate responded differently compared to the other VFAs in our experiments. When *A. taxiformis* was added to TMR incubations, net butyrate production increased from 18.7 mM to 22.3 mM.

Similarly, with ALF, net butyrate production increased from 10 mM to 13.1 mM. However, with CEL as the feed source, butyrate production was only slightly higher in positive treatments relative to negative controls. Within 24 hours, the carbon isotope composition of butyrate shifted (Figure 4). Initially, butyrate had a *δ*^13^C value of -32‰, which changed to -29‰, -30‰, and -28‰ in the first 24 hours for TMR, ALF, and CEL, respectively (Figure 4). This relatively ^13^C-enriched signal held constant for the remainder of the experiment. The hydrogen isotope composition of butyrate was consistently ^2^H-depleted when *A. taxiformis* was added to the incubation, by about 10‰ compared to the negative controls. There was no clear trend in *δ*^2^*H* with time.

## DISCUSSION

Our results demonstrate that the stable isotope compositions of VFAs in rumen fluid change systematically when a methanogen-inhibiting feed additive is introduced to rumen fluid. These differences can be explained by two shifts in the pathways of microbial fermentation: 1.) a decrease in acetate synthesis and 2.) an increase in the acrylate pathway of propionate synthesis. Both changes infer that fermentative microbial cells would receive less energy when *A. taxiformis* is added. VFA concentration profiles independently support our conclusions, though they are less informative of fermentation pathways in the rumen itself, where host consumption of VFAs prevents production rate measurements. However, we predict that VFA *δ*^13^C and *δ*^2^H measurements will translate to *in vivo* studies, quantifying the response of ruminant fermentation to metabolic inhibitors and providing important context in the design and optimization methane mitigation strategies.

### Propionate *δ*^2^H values suggest partitioning to the acrylate pathway

Propionate is an important metabolite for redox balancing the rumen microbiome (12). Fermentation of sugars must balance the production of reducing equivalents like nicotinamide adenine dinucleotide (NADH) from glycolysis with their consumption during VFA and H_2_ synthesis. Each VFA synthesis pathway uses a different number of NADH molecules (or other reducing equivalent), but propionate is the only VFA that uses more reducing equivalents than are produced during glycolysis (Equations 1-5). Propionate is thus a net sink of reducing equivalents. Acetate synthesis, on the other hand, produces two additional reducing equivalents. However, synthesizing acetate creates two additional molecules of ATP. VFA production is a tradeoff between redox balance and energy generation.

Complicating this further are the two distinct synthesis pathways for making propionate, the acrylate and succinate pathways, which influence the ATP budget of ruminant fermentation (Figure 5). The succinate pathway can generate chemiosmotic energy through a membrane-bound fumarate reductase, while the acrylate pathway does not. As such, the ratio of these two pathways determines how energetically efficient propionate synthesis can be in the rumen and has been speculated to change when methanogenesis is inhibited (32; 33). From an applied perspective, microbiome engineers hope to promote propionate synthesis to shuttle the accumulating H_2_ toward propionate production. These engineering applications would benefit from an understanding of which synthesis pathway is best adapted to the new ecology of rumen with methanogenesis inhibited. However, the acrylate and succinate pathways are indistinguishable based on VFA profiles alone.

We found that the hydrogen isotope composition of propionate had a clear trend through time in positive treatments, indicating a shift in the balance between the two propionate synthesis pathways during the inhibition of methanogenesis. In the acrylate pathway, lactate dehydrogenase (LDH) and acrylate-CoA dehydrogenase both use NADH as an electron carrier. Hydride transfer reactions from NADH to the carbon skeleton of metabolites tend to have large KIEs (e.g. LDH’s KIE >1000‰) (34). As such, the contribution of NADH-derived hydrogen atoms has been shown to influence the hydrogen isotope composition of bacterial lipids and amino acids (35; 36; 37). There are also no opportunities for either of the NADH-derived hydrogen atoms to exchange with water once they are carbon-bound in the acrylate pathway (Figure 5, red). Conversely, in the succinate pathway, only one hydrogen atom is contributed by NADH and it can be replaced via water exchange in subsequent reactions. Accounting for all these factors, we estimate that 40% vs. 15% of propionate’s hydrogen atoms are NADH-derived in the acrylate pathway vs. succinate pathway, respectively (Figure 5). Given the large KIEs associated with NADH dehydrogenase reactions, we hypothesize that propionate synthesized from acrylate would have a lower *δ*^2^H value than succinate-derived propionate. Thus, shifting metabolism into the acrylate pathway and away from the succinate pathway likely synthesized propionate with lower *δ*^2^H values, explaining our experimental results. An alternative hypothesis that could also explain the decrease in propionate *δ*^2^*H* values over time would be a change in either the water or NAD(P)H hydrogen isotope composition when *A. taxiformis* was added. However, neither acetate nor butyrate exhibit changes in *δ*^2^*H* values with time or between conditions. These products should also reflect changes in the water and NAD(P)H isotope compositions. Thus, it’s unlikely that NAD(P)H or water caused the observed signals. Notably, the acrylate pathway is less energetically efficient in terms of ATP production. While propionate synthesis is a common response to methanogen inhibitors, it may come at a higher energetic cost than previously thought, decreasing ATP yields per microbial cell and possibly the overall efficiency of fermentation in the rumen (Figure 6).

**Figure 6.**
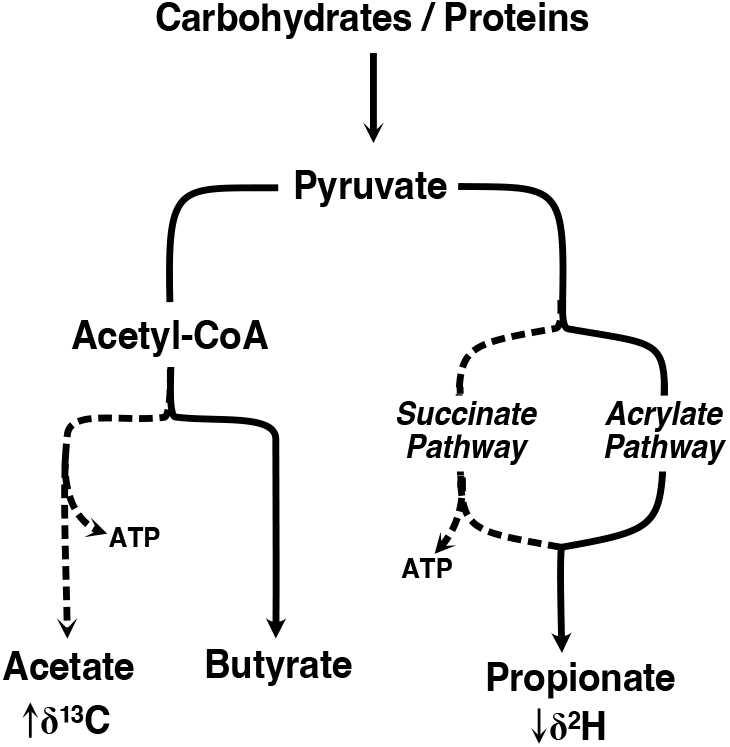
Summary of changes to microbial fermentation during methanogenesis inhibition by addition of *A. taxiformis*, highlighting that the pathways with reduced flux are ATP-producing pathways. Dotted and solid lines represent decreased and increased fluxes through those pathways, respectively. The systematic isotopic shifts observed are reported below acetate and propionate.

### Acetate *δ*^13^C values reflect shifts towards other VFAs under methanogen inhibition

Acetate is the most abundant VFA in the rumen and a critical source of energy for the animal host. As described above, acetate synthesis is a net source of reducing equivalent for the cell; however, it creates additional ATP molecules, lending cells more energy to grow and degrade plant material. Often, the rate of acetate synthesis *in vivo* is estimated from propionate-acetate ratios, which tend to increase when feed additives are administered(38; 11; 12). However, these trends lack a metabolic interpretation, as they cannot separate increased propionate synthesis from decreased acetate synthesis.

We found that the acetate was consistently ^13^C-enriched compared to the feed. Even in negative controls, acetate is 2-4‰ ^13^C-enriched, similar to isotope fractionations observed in pure cultures of fermentative bacteria (25; 22). In pure culture, ^13^C -enrichment of acetate results from other reactions that consume the precursor metabolite to acetate synthesis — acetyl-CoA. These other reactions include butyrate production, the TCA cycle, and lipid biosynthesis, each of which expresses a strong KIE (5-15‰) that leave acetyl-CoA, and subsequently acetate, ^13^C -enriched. (25). Measurements of butyrate validated this interpretation. Across all six experimental conditions, the *δ*^13^*C* value of butyrate was at least 3‰ depleted compared to the feed and 5-8‰ depleted compared to acetate, suggesting that butyrate synthesis partially drives the ^13^C - enrichment of acetate. Ni et al. (2025) (26) found a nearly identical pattern of *δ*^13^*C* values, where acetate *δ*^13^*C* values were 5-7‰ lower than those of butyrate. The remarkable agreement between two independent studies performed with different analytical techniques on distinct animal specimens points to a common metabolic mechanism, namely the acetyl-CoA node.

We hypothesized that the offset between acetate and the feed *δ*^13^C values could be a proxy for the relative rate of acetate synthesis. Specifically, it would constrain the relative amount of acetyl-CoA used to synthesize acetate, known as the branching ratio (f_acc_). As f_acc_ approaches 100%, nearly all acetyl-CoA is used for acetate synthesis, and the isotopic offset between acetate and feed should approach 0‰. As f_acc_ decreases to 0%, acetyl-CoA is entirely partitioned to generate butyrate or other metabolic end products and the offset linearly approaches the average KIE of these other enzymes consuming acetyl-CoA (See Supplementary Text for details). In this study, the offset increased from 2-3‰ to 6-8‰ between negative and positive treatments (Figure 4), indicating a decrease in f_*acc*_ when *A. taxiformis* was added. Assuming that the average KIE of the other reactions consuming acetyl-CoA is the same as butyrate synthesis (10‰), f_*acc*_ decreased 40-50% when methanogenesis was inhibited. Consistent with this estimate, net production of acetate lowered by 66%, 42%, and 52% for TMR, ALF, and CEL conditions, respectively. Meanwhile, butyrate production increased by up to 30%. Taken together, these data suggest that when methanogenesis was inhibited, acetyl-CoA was partitioned away from acetate and toward butyrate synthesis or other acetyl-CoA consumption reactions like initiating the TCA cycle (Figure S1). This represents a loss of metabolic energy, because acetate synthesis creates more ATP for the fermenting cells (Equation 3). Thus, if the same decrease in acetate synthesis were to occur *in vivo*, it would suggest that the animal host and fermenting microorganisms themselves are receiving less energy from the same amount of plant material (Figure 6).

Understanding the relative rates of VFA production in the rumen is a crucial metric for predicting animal health and guiding methane mitigation strategies. Measuring acetate production via concentration was only tractable in this study because the experiments were *in vitro* external incubations, removing consumption of the VFAs as a confounding process. During *in vivo* studies with animals given feed additives, direct production rate estimates are not possible, but the isotope compositions of VFAs provides a feasible proxy to fill this gap.

### Acetate *δ*^13^C and *δ*^2^*H* values as a proxy for acetogenesis

Acetogenesis, a metabolism that uses CO_2_ and H_2_ to generate acetate, is a promising target as an alternative metabolic H_2_ sink and as an additional source of VFA substrate to the animal host. However, acetogenesis and fermentation are diffult to distinguish with VFA concentrations alone. We initially sought to quantify the relative contributions of this metabolism using the isotope composition of acetate, since acetogenesis produces acetate with extremely low *δ*^2^*H* (<-300‰) and *δ*^13^*C* (<-50‰) values (21). We hypothesized that there would be a shift towards more negative *δ*^13^*C* and *δ*^2^*H* values in the *A. taxiformis* conditions, assuming that acetogens would take advantage of high H_2_ concentrations. However, we saw the opposite *δ*^13^*C* signal and no change in *δ*^2^*H* values, indicating that acetogens did not replace methanogens. This result is in contrast to a recent study that found a ∼2‰ depletion of acetate upon *in vitro* inhibition of methanogenesis with a different feed additive, 3-nitrooxypropanol. Ni et al. (26) inferred active acetogenesis from these data, but given that fermentation has a variable isotopic fractionation, it remained difficult to quantify the contribution of acetogenesis. The tandem proxy of *δ*^13^*C* and *δ*^2^*H* measurements, provided by Orbitrap-MS, would constrain the relative contributions of acetogenesis and fermentation, as *δ*^2^*H* values appear unaffected by the remodeling of fermentative metabolic pathways but highly sensitive to acetogenesis. Notably, the only two studies to have measured VFA isotopic properties after methanogenesis inhibition found opposite results, suggesting that acetogenesis stimulation is not a universal response to different mitigation strategies. Future studies that seek to stimulate acetogenic populations can use the isotopic methods and models developed here to quantify their success.

### Possible translation of isotopic tools to in vivo studies

Batch *in vitro* incubations of rumen fluid are not fully representative of the in vivo conditions within the animal host (38). Most notably, the host continually absorbs organic acids into their blood to be used for carbon and energy. As such, changes to the VFA concentration profile cannot be solely attributed to shifts in the ruminant fermentation. We predict that the tools and interpretations built in this study will translate to in vivo investigations. The diffusion of VFAs into the host blood stream may impart an isotope effect, which can be quantified and is unlikely to change between animals or conditions. This consumption fractionation may explain why initial acetate *δ*^13^C values are offset from the negative control time points. Thus, by correcting for the consistent isotopic fractionations of blood diffusion, we can quantify the *δ*^13^C and *δ*^2^H values of VFAs produced by fermentation, which in turn will reveal shifts in their metabolic pathways.

### ESI-Orbitrap isotomics as a tool for rumen microbiology

Here, we provide the most detailed characterization and interpretation to date of VFA *δ*^13^C and *δ*^2^H values from ruminant animals. We have enabled such analyses by developing a rapid method to determine the carbon and hydrogen isotope composition of VFA fermentation products with ESI-Oribtrap MS and coupling these data to conceptual bioisotopic models. As an analytical tool, ESI-Orbitrap isotope ratio methods were well suited for these analyses. Since acetate, propionate and butyrate are present at high concentrations, minimal preparatory work was required. Samples were filtered, diluted and directly injected onto the instrument through an autosampler. Our method uses only 5 *µ*L of rumen fluid and within 30 minutes of analysis time, all six isotopic properties (*δ*^13^C and *δ*^2^H of three VFAs) were quantified with useful precision. Furthermore, we demonstrate that isotopic measurements can differentiate the pathways of propionate synthesis, quantify the ratio of acetate-to-butyrate synthesis rates, and verify the presence of acetogenesis. Each represents an important piece of information for the continuing effort to sustainably mitigate methane emissions from ruminant animals. In the future, isotopic measurements can be integrated into quantitative isotopic models of fermentative metabolism to gain a more refined interpretation of the signals observed here (25; 39).

## CONCLUSIONS

Feed additives and other strategies that reduce methane emissions from enteric fermentation could transform livestock agriculture into a more environmentally sustainable industry. However, the implications of these additives on the microbial community that ferments plant biomass into bioavailable VFAs have not been fully elucidated. Current techniques for assessing the rates and mechanisms of microbial fermentation in the rumen cannot distinguish metabolic pathways that create the same fermentation products nor can they assess the in vivo rate of VFA generation. We present VFA isotope compositions as a novel tool to help fill this gap. ESI-Orbitrap MS methods are well designed for such a task with rapid analysis times, little-to-no sample preparation, and high biological reproducibility. The changes in VFA *δ*^13^C and *δ*^2^H values can yield new information about how fermentation is responding to the inhibition of methanogenesis. Namely, the hydrogen isotope composition of propionate reveals the importance of the less energetic acrylate synthesis pathways and the isotopic offset between acetate and cow feed points to a decrease in acetate synthesis. Both shifts suggest a loss of energetic efficiency during microbial fermentation when methanogenesis was inhibited. Our method can also distinguish acetogenesis and fermentation, though we did not observe acetogenesis in our study. Finally, we predict that stable isotope measurements will be translatable to the animal host environment, where they could help to design and optimize methane mitigation strategies.

## ACKNOWLEDGEMENTS

The authors gratefully acknowledge undergraduate assistants in the Hess Lab for their assistance in running experiments. Portions of the paper were developed from the thesis of E.P.M. Funding for this work came from an NSF Graduate Research Fellowship DGE-1745301 (to E.P.M.) and a Caltech Linde Center Grant #25570006 (to A.L.S.). We also thank the Resnick Sustainability Institute for support through an RSI explorer grant.

