## Supplementary Materials for "Shifts in ruminant fermentation during inhibition of methanogenesis are reflected in the isotope compositions of volatile fatty acids"

### Supplemental material

#### Constraining branching ratios with isotope compositions

In the main text, the carbon isotopic offset between acetate and feed is used as a proxy for the the branching ratio ($f_{acc}$) of fluxes at the acetyl-CoA node of metabolism, where $f_{acc}$ is the fraction of acetyl-CoA that is used for acetate synthesis (Figure S1). For the purposes of this initial study, it represents an overly simplified description of fermentative metabolism. However, regardless of the nuance of the model parameters or assumptions, the directionality of the proxy remains true: As $f_{acc}$ increases, the positive offset between acetate $\delta^{13}C$ and feed $\delta^{13}C$ values will increase and vice versa. Thus, even in the absence of this quantitative proxy, the isotopic enrichment of acetate holds valuable information. The following section describes the quantitative proxy mathematically.

First we define $f_{acc}$

$$f_{acc} = \frac{\phi_{acetate}}{\phi_{acetylCoA}} \tag{1}$$

Where $\phi_{acetate}$ is the flux of acetate synthesis and $\phi_{acetyl-CoA}$ is the flux of acetyl-CoA production from upstream metabolism.

By mass balance, the weighted average isotope composition of all products synthesized from acetyl-CoA should be equivalent to the isotope composition of the feed on average:

$$\delta^{13}C_{acetyl-CoA} = f_{acc}\delta^{13}C_{acetate} + (1 - f_{acc})\delta^{13}C_{other} \tag{2}$$

Where $\delta^{13}C_{other}$ represents the carbon isotope composition of all metabolic products synthesized from acetyl-CoA other than acetate and $f_{acc} + f_{other} = 1$

It was shown in a previous study of fermentative bacteria that minimal carbon isotope fractionations ($\sim$1‰) occur during the synthesis of acetyl-CoA from a starting sugar substrate, so we assume that synthesized acetyl-

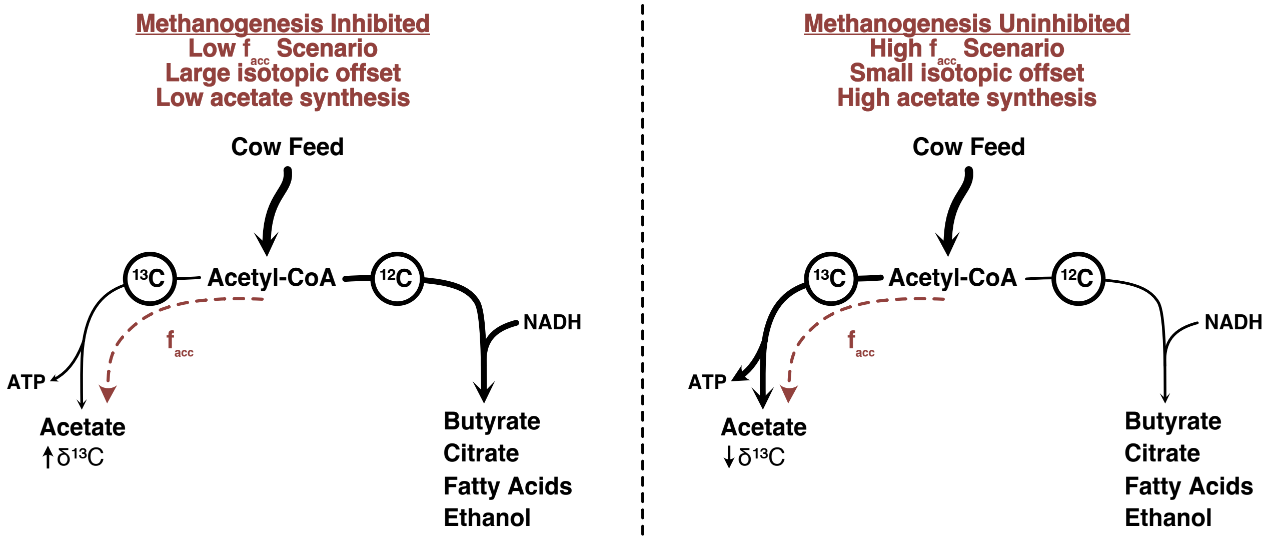


Figure 1: The acetyl-CoA branch point of metabolism constrained by the carbon isotope offset between cow feed and acetate. Negative controls were characterized by low offsets caused by high branching ratios toward acetate synthesis (right). Meanwhile, incubations where methanogenesis was inhibited were characterized by high offsets caused by low branching ratios toward acetate synthesis (left). Adapted from (1)

CoA has the same isotope composition as the feed ($\delta^{13}C_{OM}$).

$$\delta^{13}C_{OM} = f_{acc}\delta^{13}C_{acetate} + (1 - f_{acc})\delta^{13}C_{other} \tag{3}$$

The kinetic isotope effect (KIE) for acetate synthesis ($\varepsilon_{acetate}$) and other syntheses ($\varepsilon_{other}$) are expressed on a steady state pool of acetyl-CoA, which is balanced between production upstream from glycolysis and consumption by these downstream reactions. The steady state pool of acetyl-CoA has a distinct isotope composition ($\delta^{13}C_{acetyl_{C}oA,ss}$) from the "instantaneously" synthesized acetyl-CoA, which is defined by these KIEs. See (2) for more details.

$$\varepsilon_{acetate} = \delta^{13}C_{acetylCoA,ss} - \delta^{13}C_{acetate} \tag{4}$$

$$\varepsilon_{other} = \delta^{13}C_{acetylCoA,ss} - \delta^{13}C_{other} \tag{5}$$

We assume that $\varepsilon_{acetate}$ is zero, in line with previous studies (1; 3) as it involves no bond cleavage, condensation, carbon atom hybridization changes or redox changes (i.e. $\delta^{13}C_{acetate} = \delta^{13}C_{acetylCoA,ss}$). Substituting Equations S4 and S5 into Equation S3 and solving for f$_{acc}$then yields:

$$f_{acc} = 1 - \frac{\delta^{13}C_{acetate} - \delta^{13}C_{OM}}{\varepsilon_{other}} \tag{6}$$

Equation 6 demonstrates that when all of acetyl-CoA is used to make acetate ($f_{acc} = 1$), the isotopic offset is 0. As $f_{acc}$ decreases, the offset approaches $\varepsilon_{other}$, imposing the directionality of the proxy.

The reactions encompassed in $\varepsilon_{other}$ include citrate synthase (to the TCA cycle), acetyl-CoA carboxylase (to fatty acid biosynthesis), acetyl-CoA acyltransferase (to butyrate production), and alcohol dehydrogenase (to ethanol production). In previous work these enzymes were found to have similar magnitude KIEs from 5-15‰, and assume for simplicity that $\varepsilon_{other}$ is 10‰ in the main text. This assumption allows us to explicitly calculate $f_{acc}$ at a given offset value. For example, the offset increased from 2-3‰ to 6-8‰ between with and without *A. taxiformis* , respectively. Calculated $f_{acc}$ change from 0.7-0.8 and 0.2-0.4 between controls and positive treatments (Figure S1). This indicates that the branching ratio toward acetate synthesis at the acetyl-CoA node of metabolism decreased by 30-50% when methanogenesis was added, consistent with a 42-66% drop in acetate synthesis and up to 30% increase in butyrate synthesis across the various feed types (Figure 3, main text).

Table 1: HESI-Orbitrap MS Parameters

| | |
|---|---|
| **ESI Parameters** | |
| Polarity | Negative |
| Sheath Gas Flow | 10 |
| Auxiliary Gas Flow | 3 |
| Sweep Gas Flow | 3 |
| Spray Voltage | 3.1 |
| Spray Current | ¡0.2 $\mu$A |
| Auxiliary Gas Temperature | 100°C |
| Capillary Temperature | 320°C |
| **Acetate MS Parameters** | |
| Time | 0 — 11 minutes |
| Quadrupole Filter Range | 57 - 62 m/z |
| Resolution | 60,000 (at 200 m/z) |
| AGC | $1 \times 10^6$ |
| Microscans | 1 |
| S-Lens Radio Frequency Level | 60% |
| **Propionate MS Parameters** | |
| Time | 11 — 17 minutes |
| Quadrupole Filter Range | 72 - 75 m/z |
| Resolution | 60,000 (at 200 m/z) |
| AGC | $1 \times 10^6$ |
| Microscans | 1 |
| S-Lens Radio Frequency Level | 60% |
| **Butyrate MS Parameters** | |
| Time | 17 — 24 minutes |
| Quadrupole Filter Range | 85 - 90 m/z |
| Resolution | 120,000 (at 200 m/z) |
| AGC | $1 \times 10^6$ |
| Microscans | 1 |
| S-Lens Radio Frequency Level | 60% |

Table 2: Randomization of Ankom vessels and their endpoint pH measured after 72 hours. Positive and negative treatements indicate incubations with and without *A. taxiforms*, respectively. Abbreviations: CEL, cellulose; TMR, total mean ration; ALF, alfalfa

| **Vessel #** | **Feed Type** | **Treatment** | **Endpoint pH** |
|---|---|---|---|
| 1 | ALF | Positive | 5.90 |
| 2 | ALF | Negative | 5.74 |
| 3 | TMR | Negative | 5.26 |
| 4 | ALF | Negative | 5.67 |
| 5 | CEL | Negative | 5.99 |
| 6 | TMR | Positive | 5.95 |
| 7 | ALF | Negative | 5.65 |
| 8 | ALF | Positive | 5.93 |
| 9 | CEL | Negative | 6.06 |
| 10 | CEL | Negative | 5.87 |
| 11 | CEL | Positive | 6.27 |
| 12 | TMR | Positive | 5.53 |
| 13 | ALF | Negative | 5.65 |
| 14 | CEL | Negative | 5.87 |
| 15 | TMR | Positive | 5.43 |
| 16 | CEL | Positive | 6.25 |
| 17 | CEL | Positive | 6.26 |
| 18 | TMR | Negative | 5.23 |
| 19 | ALF | Positive | 6.02 |
| 20 | TMR | Negative | 5.33 |
| 21 | TMR | Negative | 5.10 |
| 22 | CEL | Positive | 6.25 |
| 23 | TMR | Positive | 6.49 |
| 24 | ALF | Positive | 5.94 |

Table 3: Gas production at 24 hour timepoint.

| **Vessel #** | **Cond** | **Total Gas (mL)** | **$CH_4$ (mL/g)** | **$CO_2$ (mL/g)** | **$CH_4$ (‰)** | **$CO_2$ (‰)** |
|---|---|---|---|---|---|---|
| 1 | ALF+ | 210 | 0.3 | 36.1 | -57.9 | -14.8 |
| 2 | ALF- | 168 | 9.7 | 14.5 | -59.7 | -11.9 |
| 3 | TMR- | 200 | 11.2 | 17.9 | -58.3 | -10.3 |
| 4 | ALF- | 180 | 12.0 | 17.8 | -61.6 | -12.2 |
| 5 | CEL- | 122 | 5.3 | 9.6 | -56.0 | -8.6 |
| 6 | TMR+ | 160 | 0.2 | 18.2 | -59.4 | -14.2 |
| 7 | ALF- | 240 | 16.9 | 31.5 | -62.5 | -12.6 |
| 8 | ALF+ | 172 | 0.5 | 25.7 | -55.7 | -15.1 |
| 9 | CEL- | 126 | 6.6 | 12.9 | -60.7 | -9.8 |
| 10 | CEL- | 150 | 10.0 | 14.7 | -59.3 | -9.2 |
| 11 | CEL+ | 105 | 0.9 | 8.6 | -54.6 | -12.4 |
| 12 | TMR+ | 240 | 0.6 | 43.5 | -55.8 | -14.8 |
| 13 | ALF- | 210 | 13.8 | 22.7 | -60.7 | -12.3 |
| 14 | CEL- | 131 | 10.6 | 15.5 | -59.4 | -9.2 |
| 15 | TMR+ | 192 | 0.4 | 52.2 | -56.4 | -13.8 |
| 16 | CEL+ | 128 | 1.1 | 17.5 | -53.3 | -13.2 |
| 17 | CEL+ | 127 | 1.2 | 13.0 | -56.9 | -13.1 |
| 18 | TMR- | 196 | 12.7 | 24.6 | -58.8 | -10.9 |
| 19 | ALF+ | 158 | 0.5 | 22.1 | -58.4 | -15.2 |
| 20 | TMR- | 182 | 11.0 | 20.1 | -58.1 | -10.8 |
| 21 | TMR- | 205 | 15.5 | 37.3 | -58.5 | -10.6 |
| 22 | CEL+ | 131 | 1.0 | 21.2 | -51.5 | -12.1 |
| 23 | TMR+ | 205 | 0.4 | 34.4 | -53.2 | -13.2 |
| 24 | ALF+ | 148 | 0.3 | 21.9 | -58.1 | -13.9 |

Table 4: Gas production at 48 hour timepoint.

| Vessel # | Cond | Total Gas (mL) | $CH_4$ (mL/g) | $CO_2$ (mL/g) | $CO_2$ (‰) | $CH_4$ (‰) |
|---|---|---|---|---|---|---|
| 1 | ALF+ | 161 | 0.0 | 29.5 | -18.4 | — |
| 2 | ALF- | 182 | 7.7 | 25.1 | -14.5 | -60.8 |
| 3 | TMR- | 175 | 9.3 | 29.6 | -13.0 | -60.6 |
| 4 | ALF- | 186 | 7.5 | 27.6 | -15.2 | -63.3 |
| 5 | CEL- | 96 | 1.8 | 12.9 | -10.0 | -65.5 |
| 6 | TMR+ | 187 | 0.0 | 36.2 | -17.5 | — |
| 7 | ALF- | 190 | 9.0 | 37.9 | -15.8 | -62.4 |
| 8 | ALF+ | 151 | 0.0 | 28.58 | -17.8 | — |
| 9 | CEL- | 92 | 1.2 | 8.9 | -9.4 | -61.5 |
| 10 | CEL- | 91 | 1.6 | 12.0 | -9.3 | -63.1 |
| 11 | CEL+ | 85 | 0.0 | 9.7 | -13.2 | — |
| 12 | TMR+ | 210 | 0.0 | 49.8 | -19.2 | — |
| 13 | ALF- | 185 | 8.5 | 34.0 | -15.1 | -62.7 |
| 14 | CEL- | 110 | 2.2 | 14.1 | -10.3 | -64.2 |
| 15 | TMR+ | 182 | 0.0 | 69.2 | -17.7 | — |
| 16 | CEL+ | 66 | 0.0 | 7.2 | -13.5 | — |
| 17 | CEL+ | 76 | 0.0 | 8.8 | -13.4 | — |
| 18 | TMR- | 180 | 8.7 | 33.4 | -13.5 | -59.0 |
| 19 | ALF+ | 168 | 0.0 | 38.3 | -18.6 | — |
| 20 | TMR- | 209 | 8.5 | 36.6 | -12.2 | -57.5 |
| 21 | TMR- | 169 | 8.5 | 38.9 | -13.4 | -58.9 |
| 22 | CEL+ | 80 | 0.0 | 11.0 | -14.0 | — |
| 23 | TMR+ | 182 | 0.0 | 48.4 | -18.0 | — |
| 24 | ALF+ | 157 | 0.1 | 34.8 | -18.6 | — |

Table 5: Gas production at 72 hour timepoint.

| Vessel # | Cond | Total Gas (mL) | $CH_4$ (mL/g) | $CO_2$ (mL/g) | $CO_2$ (‰) | $CH_4$ (‰) |
|---|---|---|---|---|---|---|
| 1 | ALF+ | 111 | 0.0 | 11.0 | -19.6 | — |
| 2 | ALF- | 134 | 3.8 | 16.2 | -16.2 | — |
| 3 | TMR- | 64 | 0.6 | 4.8 | -14.8 | — |
| 4 | ALF- | 132 | 4.2 | 20.1 | -16.8 | -64.9 |
| 5 | CEL- | 41 | 0.1 | 1.1 | -7.1 | — |
| 6 | TMR+ | 136 | 0.0 | 22.9 | -21.0 | — |
| 7 | ALF- | 125 | 3.5 | 15.1 | -17.4 | -70.4 |
| 8 | ALF+ | 131 | 0.0 | 16.8 | -20.7 | — |
| 9 | CEL- | 51 | 0.5 | 2.5 | -9.5 | -65.3 |
| 10 | CEL- | 49 | 0.3 | 2.5 | -9.1 | -63.4 |
| 11 | CEL+ | 41 | 0.0 | 1.9 | -13.1 | — |
| 12 | TMR+ | 136 | 0.0 | 20.0 | -22.1 | — |
| 13 | ALF- | 139 | 4.6 | 18.3 | -17.2 | -69.8 |
| 14 | CEL- | 52 | 0.5 | 3.3 | -9.8 | -64.2 |
| 15 | TMR+ | 136 | 0.0 | 19.8 | -21.5 | — |
| 16 | CEL+ | 45 | 0.0 | 2.5 | -14.4 | — |
| 17 | CEL+ | 47 | 0.0 | 2.3 | -14.3 | — |
| 18 | TMR- | 103 | 1.8 | 12.9 | -17.2 | -59.0 |
| 19 | ALF+ | 109 | 0.0 | 17.8 | -20.5 | — |
| 20 | TMR- | 105 | 2.0 | 14.5 | -16.4 | -55.5 |
| 21 | TMR- | 102 | 2.0 | 13.3 | -16.2 | -55.4 |
| 22 | CEL+ | 42 | 0.0 | 2.2 | -14.2 | — |
| 23 | TMR+ | 128 | 0.0 | 19.3 | -21.4 | — |
| 24 | ALF+ | 116 | 0.0 | 22.1 | -20.5 | — |

Table 6: VFA $\delta^{13}C$ and $\delta^{2}H$ values at 24 hour timepoint.

| | | $\delta^{13}C$ (‰, VPDB) | | | $\delta^{2}H$ (‰, VSMOW) | | |
|---|---|---|---|---|---|---|---|
| **Vessel #** | **Cond** | **Ac** | **Pro** | **But** | **Ac** | **Pro** | **But** |
| 1 | ALF+ | -20.7 | -28.1 | -32.1 | -197 | -231 | -225 |
| 2 | ALF- | -23.0 | -26.4 | -28.9 | -203 | -214 | -212 |
| 3 | TMR- | -22.2 | -25.6 | -30.1 | -208 | -217 | -226 |
| 4 | ALF- | -23.9 | -26.9 | -30.8 | -203 | -214 | -214 |
| 5 | CEL- | -21.6 | -26.2 | -28.6 | -195 | -214 | -217 |
| 6 | TMR+ | -20.7 | -28.4 | -31.1 | -192 | -224 | -235 |
| 7 | ALF- | -23.7 | -24.9 | -29.3 | -206 | -212 | -211 |
| 8 | ALF+ | -21.0 | -27.0 | -30.6 | -198 | -219 | -231 |
| 9 | CEL- | -22.4 | -25.3 | -29.3 | -209 | -201 | -215 |
| 10 | CEL- | -22.1 | -26.5 | -27.8 | -200 | -216 | -208 |
| 11 | CEL+ | -19.4 | -26.8 | -28.3 | -197 | -224 | -218 |
| 12 | TMR+ | -18.7 | -27.8 | -30.6 | -196 | -228 | -231 |
| 13 | ALF- | -24.3 | -25.0 | -28.8 | -203 | -207 | -214 |
| 14 | CEL- | -22.4 | -26.0 | -29.1 | -195 | -201 | -219 |
| 15 | TMR+ | -19.7 | -26.9 | -28.9 | -190 | -233 | -232 |
| 16 | CEL+ | -19.3 | -27.7 | -28.4 | -188 | -227 | -222 |
| 17 | CEL+ | -20.2 | -27.7 | -28.3 | -194 | -226 | -224 |
| 18 | TMR- | -23.4 | -25.7 | -31.1 | -205 | -209 | -215 |
| 19 | ALF+ | -20.2 | -26.5 | -29.5 | -203 | -217 | -226 |
| 20 | TMR- | -22.5 | -24.9 | -28.7 | -201 | -213 | -220 |
| 21 | TMR- | -23.2 | -25.1 | -30.1 | -205 | -202 | -223 |
| 22 | CEL+ | -18.9 | -26.1 | -28.7 | -197 | -215 | -217 |
| 23 | TMR+ | -19.7 | -25.0 | -28.2 | -203 | -215 | -230 |
| 24 | ALF+ | -21.2 | -26.8 | -30.5 | -196 | -224 | -225 |

Table 7: VFA $\delta^{13}C$ and $\delta^{2}H$ values at 48 hour timepoint.

| | | $\delta^{13}C$ (‰, VPDB) | | | $\delta^{2}H$ (‰, VSMOW) | | |
|---|---|---|---|---|---|---|---|
| **Vessel #** | **Cond** | **Ac** | **Pro** | **But** | **Ac** | **Pro** | **But** |
| 1 | ALF+ | -22.0 | -26.8 | -31.5 | -204 | -233 | -224 |
| 2 | ALF- | -25.1 | -26.1 | -28.3 | -207 | -214 | -211 |
| 3 | TMR- | -22.1 | -25.8 | -30.0 | -206 | -212 | -227 |
| 4 | ALF- | -24.4 | -26.5 | -29.4 | -209 | -210 | -213 |
| 5 | CEL- | -23.2 | -25.3 | -27.3 | -199 | -210 | -203 |
| 6 | TMR+ | -19.1 | -26.3 | -29.1 | -185 | -233 | -225 |
| 7 | ALF- | -26.9 | -27.1 | -29.5 | -207 | -212 | -199 |
| 8 | ALF+ | -23.2 | -28.5 | -29.1 | -195 | -227 | -211 |
| 9 | CEL- | -24.6 | -27.2 | -27.8 | -200 | -200 | -208 |
| 10 | CEL- | -25.1 | -27.2 | -26.6 | -197 | -200 | -200 |
| 11 | CEL+ | -23.3 | -28.0 | -28.6 | -187 | -222 | -214 |
| 12 | TMR+ | -19.6 | -26.9 | -28.5 | -181 | -221 | -228 |
| 13 | ALF- | -28.3 | -27.3 | -29.7 | -202 | -201 | -200 |
| 14 | CEL- | -25.3 | -25.5 | -28.0 | -203 | -198 | -205 |
| 15 | TMR+ | -17.9 | -26.4 | -28.6 | -186 | -229 | -239 |
| 16 | CEL+ | -20.6 | -26.2 | -28.4 | -201 | -220 | -212 |
| 17 | CEL+ | -19.4 | -26.4 | -28.9 | -198 | -222 | -213 |
| 18 | TMR- | -23.73 | -26.5 | -30.0 | -202 | -212 | -226 |
| 19 | ALF+ | -20.7 | -27.4 | -29.4 | -205 | -226 | -220 |
| 20 | TMR- | -24.1 | -24.5 | -30.0 | -208 | -205 | -207 |
| 21 | TMR- | -25.6 | -24.0 | -29.3 | -201 | -219 | -221 |
| 22 | CEL+ | -17.9 | -28.3 | -27.5 | -194 | -218 | -208 |
| 23 | TMR+ | -18.4 | -26.4 | -28.6 | -188 | -248 | -236 |
| 24 | ALF+ | -23.4 | -26.6 | -31.3 | -200 | -215 | -214 |

Table 8: VFA $\delta^{13}C$ and $\delta^{2}H$ values at 72 hour timepoint.

| | | $\delta^{13}C$ (‰, VPDB) | | | $\delta^{2}H$ (‰, VSMOW) | | |
|---|---|---|---|---|---|---|---|
| **Vessel #** | **Cond** | **Ac** | **Pro** | **But** | **Ac** | **Pro** | **But** |
| 1 | ALF+ | -21.0 | -27.8 | -30.2 | -201 | -251 | -227 |
| 2 | ALF- | -25.7 | -27.7 | -30.0 | -206 | -207 | -209 |
| 3 | TMR- | -21.6 | -24.6 | -30.5 | -208 | -216 | -239 |
| 4 | ALF- | -25.9 | -27.4 | -31.7 | -204 | -206 | -211 |
| 5 | CEL- | -23.2 | -26.7 | -28.7 | -205 | -211 | -203 |
| 6 | TMR+ | -11.3 | -23.7 | -29.0 | -174 | -240 | -251 |
| 7 | ALF- | -26.4 | -26.7 | -31.4 | -208 | -213 | -218 |
| 8 | ALF+ | -20.1 | -27.5 | -31.5 | -202 | -227 | -221 |
| 9 | CEL- | -22.8 | -26.5 | -29.4 | -203 | -206 | -205 |
| 10 | CEL- | -23.1 | -25.8 | -28.3 | -208 | -215 | -197 |
| 11 | CEL+ | -20.6 | -25.9 | -28.4 | -200 | -224 | -210 |
| 12 | TMR+ | -12.5 | -24.9 | -29.7 | -188 | -239 | -234 |
| 13 | ALF- | -25.0 | -27.8 | -32.4 | -211 | -215 | -201 |
| 14 | CEL- | -22.4 | -25.9 | -29.1 | -198 | -203 | -200 |
| 15 | TMR+ | -11.7 | -25.6 | -29.3 | -191 | -243 | -239 |
| 16 | CEL+ | -19.2 | -27.0 | -27.1 | -193 | -223 | -210 |
| 17 | CEL+ | -19.0 | -26.8 | -27.9 | -201 | -224 | -214 |
| 18 | TMR- | -21.4 | -26.6 | -30.8 | -208 | -223 | -230 |
| 19 | ALF+ | -22.3 | -28.1 | -33.1 | -204 | -243 | -223 |
| 20 | TMR- | -20.7 | -25.0 | -29.1 | -208 | -207 | -229 |
| 21 | TMR- | -18.4 | -25.2 | -28.6 | -208 | -216 | -227 |
| 22 | CEL+ | -18.2 | -27.6 | -26.4 | -206 | -219 | -213 |
| 23 | TMR+ | -13.4 | -24.1 | -30.0 | -186 | -245 | -236 |
| 24 | ALF+ | -20.9 | -27.3 | -30.8 | -203 | -240 | -220 |
